# History-dependent spiking facilitates efficient encoding of polarization angles in neurons of the central complex

**DOI:** 10.1101/2024.07.25.605186

**Authors:** Lisa Rother, Anna Stöckl, Keram Pfeiffer

## Abstract

Many insects use the polarization pattern of the sky for spatial orientation. Since flying insects perform rapid maneuvers, including saccadic yaw turns which alternate with translational flight, they perceive highly dynamic polarization input to their navigation system. The tuning of compass-neurons in the central complex of insects, however, has been mostly investigated with polarized-light stimuli that rotated at slow and constant velocities, and thus were lacking these natural dynamics. Here we investigated the dynamic response properties of compass-neurons, using intracellular recordings in the central complex of bumblebees. We generated naturalistic stimuli by rotating a polarizer either according to a sequence of head orientations that have been reported from freely flying bumblebees, or at constant velocities between 30°/s and 1920°/s, spanning almost the entire range of naturally occurring rotation velocities. We found that compass neurons responded reliably across the entire range of the presented stimuli. In their responses, we observed a dependency on spiking history. We further investigated this dependency using a rate code model taking spiking history into account. Extending the model to a neuronal population with different polarization tuning, which mirrored the neuronal architecture of the central complex, suggests that spiking history has a directly impact on the overall population activity, which has two effects: First, it facilitates faster responses to stimulus changes during highly dynamic flight maneuvers, and increases sensitivity for course deviations during straight flight. Second, population activity during phases of constant polarization input is reduced, which might conserve energy during straight flight.

**Significance Statement:** Many insects use the pattern of polarized light, which arises from scattering of sunlight, for spatial orientation. Neurons in the central complex, a brain structure, which governs spatial orientation, encode the orientation of the insect’s head with respect to polarized light. To investigate the dynamic properties of these neurons, we recorded intracellularly in the central complex of bumblebees and stimulated with naturalistic polarized-light stimuli. We found that neuronal activity was not only dictated by the angle of polarization, but also by the amount of previous activity. Using a modelling approach we show, that this dependency on spiking history can facilitate faster adjustment of the heading signal and leads to reduced overall activity and therefore reduced energy consumption during straight flight.

## Introduction

For accurate spatial orientation many insects use a polarized light-based compass (desert ant: Wehner, 2003, dung beetle: Birukow, 1953; Dacke et al., 2003; el Jundi et al., 2015, monarch butterfly: Heinze and Reppert, 2011, locust: Heinze and Homberg, 2009; Homberg et al., 2022, cricket: Labhart, 1988). Remarkable progress has been made over the last decades in understanding the use and the neuronal processing of the sky’s polarization pattern in insects (Wehner and Labhart, 2006; Srinivasan, 2011; Hardcastle et al., 2021). Polarization information, emanating from scattered sunlight in the atmosphere, is processed in the sky-compass pathway, which receives light information from specialized dorsally directed, polarization-sensitive photoreceptors in the dorsal part of the compound eye (dorsal rim area: DRA, Labhart, 1980; Wehner and Strasser, 1985; Labhart and Meyer, 1999; Wehner and Labhart, 2006) and relays it to neurons in the medulla, the anterior optic tubercle and eventually to the core compass network in the central complex (CX, Müller et al., 1997; Blum and Labhart, 2000; Vitzthum et al., 2002; Homberg et al., 2003; Pfeiffer et al., 2005; Heinze and Homberg, 2007; Pfeiffer and Kinoshita, 2012; Zeller et al., 2015; Held et al., 2016; Sayre et al., 2021). The CX is a group of midline-spanning neuropils of the insect brain and serves as an internal compass in bees and other insects including flies, locusts, crickets, butterflies, and beetles (Heinze, 2014; Pfeiffer, 2023). The network of CX neurons encodes an internal representation of the animal’s heading direction relative to its environment (Collett and Collett, 2002; Seelig and Jayaraman, 2015; Kim et al., 2017). This neuronal representation of heading direction must be constantly updated during active locomotion. Studies in a variety of insect species, targeting the physiological properties of polarization sensitive CX neurons, have been using simplified stimuli. Usually, a backlit polarizer was presented in the dorsal visual field of the animal. The polarizer was then rotated at a constant angular velocity (usually between 30°/s and 90°/s) to present the different angles of polarization (AoP) that are present in the polarization pattern of the sky (e.g. Labhart, 1988; Pfeiffer et al., 2005; Pegel et al., 2019). These stimuli are suitable to probe the overall response properties and the angular tuning of the neurons, but they teach us very little about the neurons’ dynamic properties and their coding properties in response to naturalistic input as it occurs during flight. Previous efforts towards gaining a better understanding of what polarization-sensitive neurons might signal under natural circumstances included presentation of different degrees of polarization (Pfeiffer et al., 2011; Hensgen et al., 2022), or successively stimulating the entire receptive fields of the neurons (Bech et al., 2014; Zittrell et al., 2020) but no attempts have been made to explore the influence of the temporal properties of the stimulus. The flight trajectory of many insect species is composed of straight segments combined with fast turns about their vertical body axis (Riabinina et al., 2014; Boeddeker et al., 2015). For the visual system, these rapid changes of gaze direction are analogous to the human eye movements and are therefore called saccades (Collett and Land, 1975). Saccades can reach peak velocities ranging from below 250°/s to 2000°/s (Riabinina et al., 2014; Boeddeker et al., 2015). When animals move through the world, they experience a continuously changing retinal image which leads to a highly dynamic input into the visual system, with spatiotemporal properties that differ fundamentally from those of the sky-compass stimuli used in previous studies.

To explore the dynamic properties of CX neurons, we performed intracellular recordings in compass neurons of the CX of *B. terrestris*, using naturalistic stimuli simulating the perception of free-flying bumblebees. To better understand the measured responses, we designed a rate-code model of neuronal activity. The model calculations show that spiking history is an important factor that shapes the dynamic response properties of central-complex neurons.

## Material and Methods

### Animals and preparation

Bumblebee (*Bombus terrestris*) colonies were obtained from two commercial suppliers (Biobest NV, Belgium and Koppert B.V., Berkel en Rodenrijs, NL). Animals were kept in two-chambered wooden boxes in a climate chamber at 25 °C and 55% rH under “white” light (12:12 LD) and provisioned *ad-libitum* food. For experiments, the animals were immobilized in the freezer for 5–20 min, then mounted onto a custom-built metal holder using dental wax (Omnident, Rodgau, Germany). A window was cut frontally into the head capsule and the gut, tracheal air sacs, glands and muscles in the head capsule were carefully removed to get access to the brain and to reduce movements. Finally, the neural sheath of the brain was partly removed using fine tweezers, to allow penetration of the recording electrode. The brain was kept submerged in bee ringer (in mM: NaCl, 130; KCl, 5; MgCl_2_, 4; CaCl_2_, 5; HEPES, 15; Glucose, 25; Sucrose, 160) during preparation and recording.

### Intracellular Recordings

We measured neural activity with sharp microelectrodes drawn from omega-shaped borosilicate glass (inside diameter: 0.75 mm, outside diameter: 1.5 mm, Hilgenberg, Malsfeld, Germany), with a Flaming/Brown filament puller (P-97, Sutter Instrument, Novato, CA, USA). The electrode tips were filled with 4% Neurobiotin tracer (Vector Laboratories; Burlingame, CA, USA) in 1 M KCl and backed up with 1 M KCl. Signals were amplified 10 × with a BA-03X amplifier (npi electronic, Tamm, Germany), visualized with a digital oscilloscope (Hameg Instruments GmbH, Frankfurt, Germany) and made audible with a custom-built audio monitor. Data were low pass filtered at 2.5 kHz, digitized with a Power1401-mkII (Cambridge Electronic Design, Cambridge, UK) at a sampling rate of 20 kHz and stored on a PC using Spike2 (Cambridge Electronic Design, Cambridge, UK). After the recording, Neurobiotin was iontophoretically injected into the neuron by applying a positive current of 1.5 nA for 1 – 3 minutes.

### Histology and Image Acquisition

Brains were fixed in Neurobiotin-Fixative (4% Paraformaldehyde (PFA), 0.2% saturated picric acid and 0.25% glutaraldehyde in 0.4 g/l NaH_2_PO_4_ * H_2_O, 0.815 g/l NaH_2_PO_4_ * 2H_2_O,) overnight at 4 °C and afterwards dissected. Following rinses (4 × 15 min) in PBS (phosphate buffered saline, containing (in mM): NaCl, 137; KCl, 2.7; Na_2_HPO_4_, 8; KH_2_PO_4_, 1.4) brains were incubated with Alexa568-conjugated Streptavidin (1:1000 in PBS 0.1 M plus 0.5% Triton X-100 (PBT)) at 4 °C for three days. After incubation, the brains were rinsed in PBT (2 x 30 min) and PBS (3 x 30 min). Brains were dehydrated in an increasing ethanol series (30%, 50%, 70%, 90%, 95%, 100%, 100% 15 min each) and cleared in a 1:1 mixture of 100% ethanol and methyl salicylate (Carl Roth GmbH + Co. KG, Karlsruhe, Germany) for 20 min and in pure methyl salicylate overnight. Finally, the brains were covered in Permount (Fisher Scientific, Schwerte, Germany) between two cover slips. Twelve hole-reinforcement rings (Avery Zweckform, Oberlaindern, Germany), stacked on top of each other were used as spacers.

Samples were scanned with a confocal laser scanning microscope (Leica, TCS SP8, Leica Microsystems, Wetzlar, Germany) with a 10 x water-immersion (HCX POL) or 20 × multi-immersion objective (HC PL APO 20 ×/0.75 Imm Corr CS2, Leica) with either water or oil as immersion medium.

### Stimulation

Experiments were carried out in the dark. Dorsal polarized-light stimuli were generated by passing the light of a UV-LED (Nichia NVSU233A-D1, peak wavelength 365 nm, intensity: 10^13^ photons cm^-2^ s^-1^, visual angle: 10°) through a polarizer (BVO UV, Bolder Vision Optik Inc., Boulder, CO, USA) that was mounted to a hollow shaft stepper motor (M013022, NEMA 23, Koco Motion GmbH, Dauchingen, Germany). The stepper motor was powered through a digital stepping drive (DM442, Leadshine Technology Co., China) that was controlled by an Arduino Mega 2560 (Funduino, Germany) using custom written code. The angle of polarization was monitored using a custom-built polarimeter, located behind the animal.

For the experiments, the polarizer was either rotated clockwise (cw, 0°-180°) or counter-clockwise (ccw, 180°-0°) in an ascending sequence of constant angular velocities (30°/s, 60°/s, 120°/s, 240°/s, 480°/s, 960°/s, 1440°/s, 1920°/s; 12 s each) or using a naturalistic stimulus sequence derived from head yaw angles that were measured in freely flying bumblebees by Boeddeker et al. (2015). The naturalistic stimulus sequence was repeated 10-15 times for each recording. To test for spiking-history-effects after stimulation with the preferred and anti-preferred angle of polarization, we first rotated the polarizer by 360° at a velocity of 60° cw and ccw. Using a custom-written spike2 script we calculated the preferred and anti-preferred angles of polarization on-line. The polarizer was then first turned to the preferred AoP, which was presented for 10 s, followed by a 10 s period of darkness. Afterwards the polarizer was rotated to the anti-preferred AoP, the light was turned on for 10 s, followed by a 10 s period of darkness.

### Preprocessing of physiological data

Recordings were visualized and preprocessed using Spike2 (Cambridge Electronic Design, Cambridge, UK). Times of action potentials were extracted based on crossing of a manually set threshold using a custom written script. Raster plots and peristimulus time histograms (PSTHs) were created using the Stimulus Histogram function with a bin size of 0.01 s. All data were exported as .mat files and all subsequent analyses were performed using custom written scripts in MATLAB (version 2020b, The MathWorks).

### Calculating preferred AoP

Every action potential during each 360° rotation was assigned the corresponding orientation of the polarizer at the time of the action potential. The preferred AoP (Φ_max_) was then calculated as the average of these angles. Circular histograms for responses at each rotation velocity and rotation direction were created with the CircHist package (Zittrell, 2020). Circular statistics (average angle, 95 % confidence interval, resultant vector length, Rayleigh test of uniformity and circular-linear correlation) were calculated using the Circular Statistics Toolbox (Berens, 2009).

### Circular linear correlation

To quantify the dependency between the preferred angle of polarization and the rotation velocity, we calculated the circular-linear regression between the two variables (Kempter et al., 2012). To this end we separated the neuronal responses to each rotation velocity into a first and second half of 6 s each, calculated the preferred angle of polarization for each segment and plotted it against the corresponding rotation velocity. For both cw and ccw rotation the circular-linear regression was calculated for the five highest rotation velocities. The absolute value of the slope of the regression line gave the delay of the system.

### Mean spike rate

Mean spike rate was calculated for each rotation velocity as the number of spikes divided by the duration of stimulation. It was calculated and plotted separately for the two rotation directions (clockwise and counter-clockwise).

### Computational Model

To dissect the components underlying the neurons’ responses to naturalistic stimuli, we designed a rate-code model of neuronal activity. The neuronal response (R) over time (t) to the angle of polarization (AoP) was implemented as a cos^2^-function:

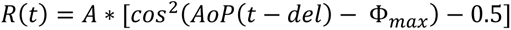

where *A* is the response amplitude, *del* is the system’s delay and *Φ_max_* is the preferred angle of polarization of the neuron. The subtraction of 0.5 accounts for polarization-opponency, i.e. the neuron is excited at Φ_max_ and inhibited at Φ_max_ – 90°.

Both during naturalistic stimulation and during stimulation with stationary polarization stimuli, we noticed a dependency of neuronal activity on spiking history. This effect led to increased activity after inhibition and to decreased activity after excitation. To accommodate for this observation, spiking history (‘his’) effects were implemented into our model as

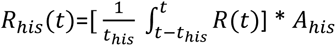

where *t_his_*is the length of a rectangular window preceding the current stimulus over which past responses are averaged and *A_his_* is a scaling factor for the spiking-history-effect.

The total response rate was then calculated as:

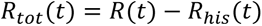

As our model was intended to represent spike rates, negative values were replaced by zeros. To compare the model to our measured neuronal data, we used PSTHs from neuronal responses to 7-15 repetitions of the naturalistic stimulus. The PSTH data were smoothed over 5 bins using the smooth function in Matlab and resampled at 1 kHz. We then used fminsearchbnd (D’Errico, 2023) in Matlab to minimise the sum of squared errors between the measured response and the model. The parameters *Φ_max_* (preferred AoP), *del* (delay), *A* (response amplitude), *t_his_* (length of history window) and *A_his_* (amplitude history response) were used as free parameters within the following boundaries: *Φ_max_*, 0° –180°; *del*, 0 s –0.3 s; *A*, 0 – 200; *t_his_*, 0 s –1.5 s; *A_his_*, 0 –1. To avoid the convergence of the fit to local minima, we repeated the fitting procedure 50 times for each of the recordings and initialized the fitting parameters each time with random start values. The fitted parameter values were then obtained from the best of the 50 fits. To assess the quality of the fit, we implemented a bootstrapping method. We repeated the fitting procedure 1000 times, but instead of fitting the model to the original PSTHs, we randomly permuted the bins of the PSTHs before the fit. For each permutation we calculated the sum of squared errors between the PSTH and the fit resulting in a distribution of sums of squared errors (Supplemental Figure 2). We then compared the sum of squared errors obtained from the fit to the original data to the sum of squared errors of the permuted data. We considered the fitted model objectively good, if the sum of squared errors of the original data was smaller than that of the lowest 5% of the permuted data. This was the case for 18 out of 24 recordings. Only these recordings were considered for further analyses. To check if the model that included spiking history indeed represented the measured data better than the model without spiking history, we repeated the fitting procedure for each neuron without the parameters *t_his_*, and *A_his_*. By eliminating spiking history from the model, we also reduced the number of model parameters from five to three. To account for this difference in parameters, we calculated the Akaike information criterion (AIC) to compare the different model candidates. This is based on the value of the log-likelihood, which is greater the better the model explains the dependent variable. In order to not classify more complex models as consistently better, the number of estimated parameters is also included as a penalty term in addition to the log-likelihood.

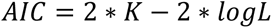

where *K* is the number of model parameters and *logL* is the log-likelihood

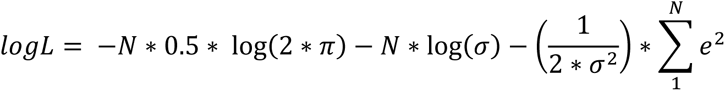

where N is the number of data points, *e* are the residuals (i.e. the difference between the data and the model) and σ is the standard deviation of the residuals.

The preferred model is the one with the lowest AIC value. AIC thus rewards the accuracy of fit of a model on the one hand, but penalizes an excessively high number of parameters.

### Population analysis

Neurons in the CX usually exist as isomorphic populations. Directional information is then jointly coded through the activity of such a population. We therefore extended our model to represent a population of CX neurons. To create a population of “standardized” neurons, we calculated the median values of the fitted parameters from the recorded TL3 neurons, except for the preferred AoP. This parameter was systematically changed from 0° to 179° in 1° increments throughout the population of neurons. We then calculated the response of this neuronal population to a 70 s series of naturalistic stimuli obtained from Boedekker et al. (2015), as well as to a series of step stimuli.

## Results

To characterize the compass neurons’ dynamic response properties, we recorded intracellularly from 37 polarization sensitive neurons (tangential neurons of the lower division of the central body: TL3=15, TL2=2, unidentified subtypes=5; columnar neurons=3; unknown=12) in the bumblebee central complex (CX). We stimulated the neurons with UV-light that was passed through a polarizer which was rotated either clockwise or counter-clockwise at velocities ranging from 30°/s to 1920°/s. Additionally, we used a naturalistic stimulus sequence generated from head orientations recorded from freely flying bumblebees (Figure 1; Boeddeker et al., 2015).

**Figure 1:**
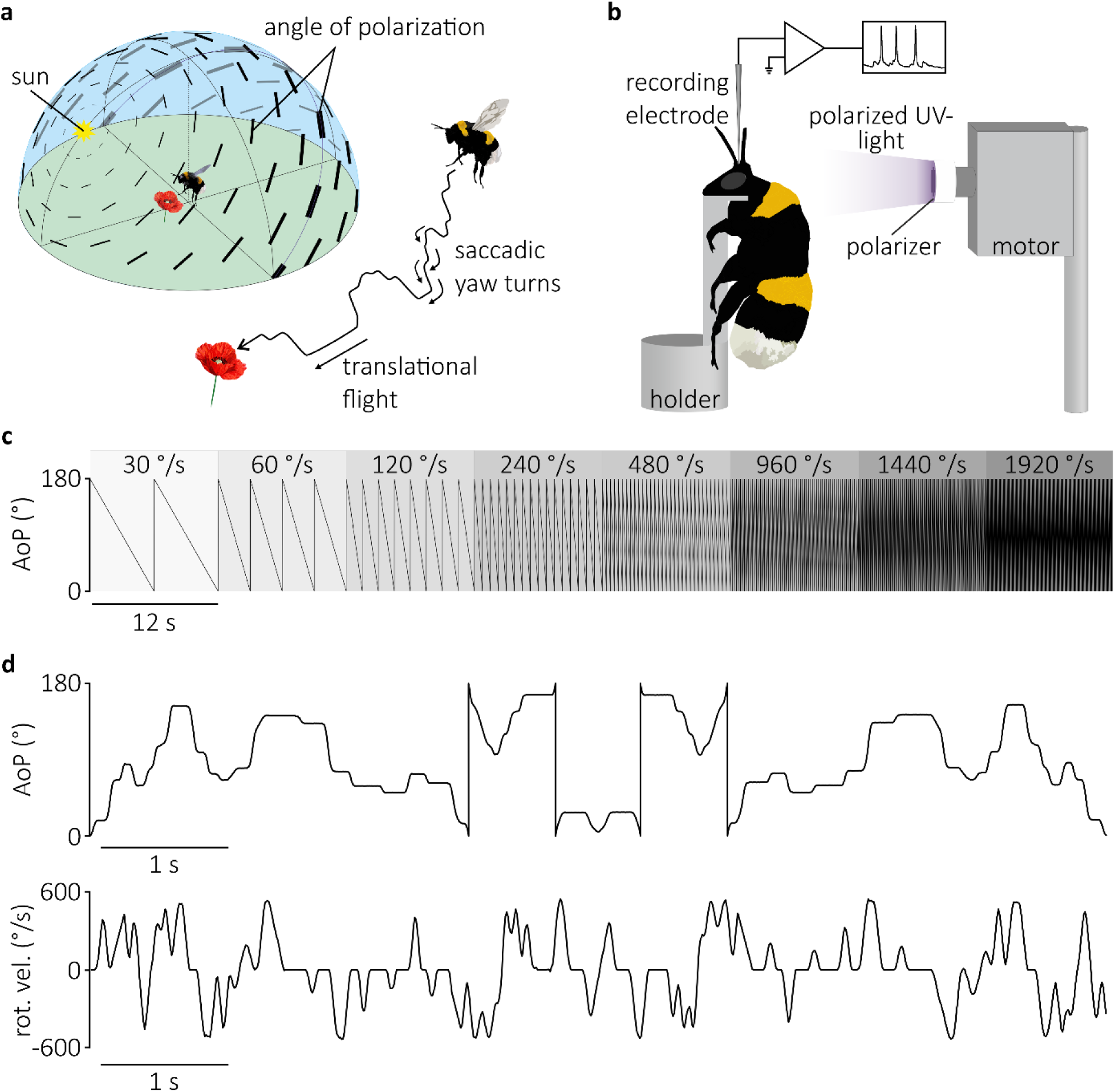
Illustration of research question, experimental setup, and stimuli. A) Bumblebees experience rapid changes in the angle of polarization as they fly under natural skies. Left: polarization pattern of the sky. Angles of polarization are oriented along concentric circles around the sun. While direct sunlight is unpolarized, the degree of polarization, indicated by the thickness of the bars, increases with increasing angular distance from the sun. Right: naturalistic flight trace with saccadic yaw turns alternating with translational flight. B) Schematic illustration of intracellular recordings from compass neurons in the central complex of *Bombus terrestris* during stimulation with a rotatable polarizer backlit by a UV-LED (365 nm). C) Counter-clockwise rotation (180°*–*0°) of the polarizer at discrete angular velocities between 30°/s and 1920 °/s. Each rotation velocity was presented for 12 s. D) Upper panel: naturalistic stimulus sequence reproducing the dynamics of an actual flight (original flight data from Boeddeker et al., 2015). Lower panel: rotation velocities of the naturalistic stimulus used in this study.

Most recorded neurons (n=15) were TL3 neurons (Figure 2a), which received input in the medial bulb and projected to the lower division of the central body (CBL). During stimulation with a continuously rotating polarizer, TL3 neurons, like most other polarization sensitive neurons, exhibited spiking activities that roughly followed a sinusoidal function with a periodicity of 180°, alternating between excitation and inhibition as shown in the example recording trace (Figure 2b). The maxima (preferred angle of polarization (AoP)) and minima (anti-preferred AoP) were separated by 90°. These typical sinusoidal responses could be observed at lower rotation velocities (Figure 2b; 30°/s and 120°/s). At 480°/s, the sinusoidal response characteristics started to break down, and were not present at 1440°/s anymore. However, while this neuron did not fire action potentials during each individual rotation of the polarizer, the spikes that were generated were significantly phase locked to the stimulus angle as tested by circular-linear correlation (Figure 2c). This means that for each rotation velocity the majority of the spikes occurred around a specific AoP.

**Figure 2:**
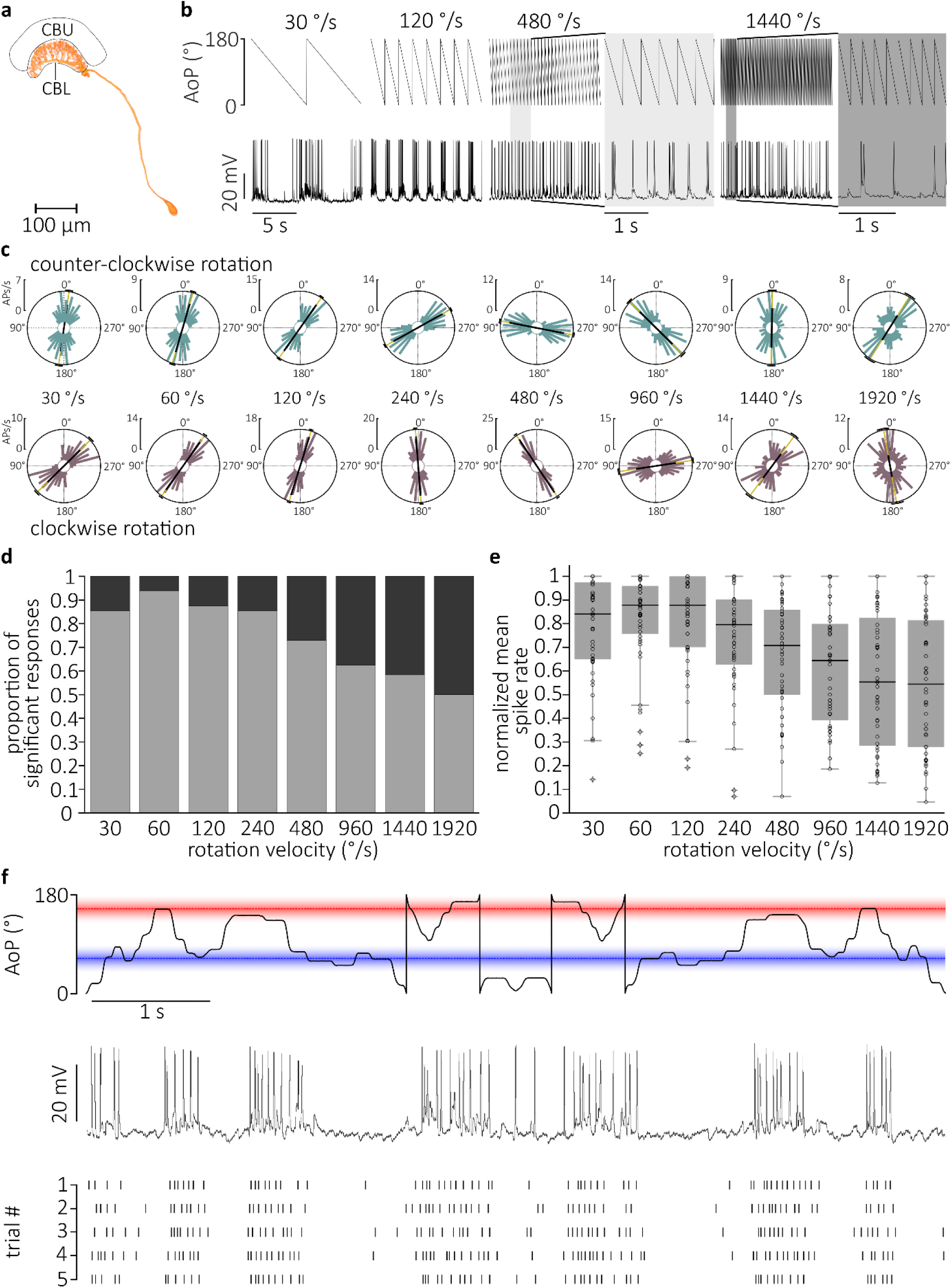
CX neurons can reliably encode naturalistic stimuli. A) Direct volume rendering of dye-injected TL3 neuron. B) Neuronal responses of TL3 neuron to selected rotation velocities (30°/s, 120°/s, 480°/s, 1440°/s) of a counter-clockwise rotation stimulus. Grey areas show responses to 480°/s and 1440°/s at a stretched timescale. C) Circular response histograms for clockwise (cw; purple) and counter-clockwise (ccw; turquoise) rotation for all presented velocities showing the preferred firing directions of the tested TL3 neuron. The mean preferred firing directions and the mean vector length are indicated by the golden/black lines. The 95% confidence intervals are given by the black arcs. D) Proportion of significantly directed responses for all recordings at the different rotation velocities (cw and ccw plotted together; n=24, N=48). E) Normalized mean spike rate for all recordings at the different rotation velocities (cw and ccw plotted together; n=24, N=48). Boxplots show median, interquartile range (IQR), whiskers with 1.5 x IQR and outliers greater than 1.5 x IQR. F) Upper panel: Naturalistic stimulus sequence according to head orientations obtained from freely flying bumblebees (Boedekker et al., 2015). Red line shows the preferred angle of polarization (AoP), the blue line the anti-preferred AoP of this neuron. Middle panel: Intracellular recording trace of the same neuron as above responding to the naturalistic stimulus sequence. Lower panel: Raster plots of five consecutive trials of stimulation with the naturalistic stimulus sequence.

Analyzing all recordings (n=24, with clockwise (cw) and counter-clockwise (ccw) rotation directions taken together: N=48) showed that depending on rotation velocity between 50% and 93.8% of the responses were significantly phase-locked to the stimulus (Figure 2d; 30°/s: 85.4%, 60°/s: 93.75%, 120°/s: 87.5%, 240°/s: 85.4%, 480°/s: 72.9%, 960°/s: 62.5%, 1440°/s: 58.3%, 1920°/s: 50%). However, the preferred AoP, differed for the different rotation velocities, and between rotation directions (Figure 2c). The changes in preferred AoP were systematic, such that the faster the polarizer was rotated the further the preferred AoP was shifted to higher angular values for cw (0°-180°) rotation or lower angular values for ccw (180°-0°) rotation (Figure 2c).

The average spiking rate during stimulation with a constantly rotating polarizer was correlated with the rotation velocity. It peaked at a rotation velocity of 60°/s and continuously decreased to either side of the peak with the lowest average activities occurring at the highest rotation velocity (Figure 2e).

The experiments with different rotation velocities showed that the CX-neurons responded with spiking patterns, that were significantly phase-locked to the stimulus and that they were capable of responding to very high rotation velocities. We next asked how the response properties we observed during stimulation with constant rotation velocities translate to a more naturalistic stimulus situation. To test this, we used a naturalistic stimulus based on a sequence of head yaw angles from a bumblebee flight (Boeddeker et al., 2015). This sequence included saccadic yaw turns and translational flight sections (Figure 2f) to simulate the temporal structure of polarization angles a bumblebee would experience while flying under the polarized sky.

Despite the rapid changes in both AoP and rotation velocity, the neurons showed very clear responses to the stimulus, which were consistent during repeated presentation of the same stimulus sequence (Figure 2f). We also noticed that during presentation of the naturalistic stimulus, identical AoPs during different parts of the stimulus lead to different levels of activity (Figure 2f; red line for preferred AoP and blue line for anti-preferred AoP), suggesting that spiking activity did not only depend on the current stimulus, but also on the spiking history.

To investigate the influence of spiking history, we analyzed the relationship between the preferred AoP and the rotation velocity of the continuously rotating stimulus. To this end, we calculated circular linear regressions (Kempter et al. 2012) between the higher rotation velocities (240°/s-1920°/s) and the corresponding preferred AoPs (separately for cw and ccw rotation; Figure 3a). The circular linear correlation analysis showed that at high rotation velocities, activity was systematically phase-delayed (indicated by the black regression line). This phase-delay was exclusively dependent on the time-delay between the stimulus and the encoding of the stimulus in the central complex. The time delay was equivalent to the absolute value of the slope of the regression line. If the delay of the system was the only factor shaping the neuronal responses, one would expect that the lower the rotation velocity, the more similar the preferred AoP for cw and ccw rotation would get, i.e. at very slow rotation velocities the preferred AoP for cw and ccw rotation should be identical. This was, however, not what we observed. Instead, we found that at low rotation velocities (30°/s-120°/s), the phase of the firing was advanced as if the delay of the system had become negative (Figure 3a). This suggests that a second mechanism, must have contributed to shaping the neuronal responses. The effect of this mechanism was directed in the opposite direction as the delay and even overrode the delay at low stimulus velocities, effectively creating a phase-advancement.

**Figure 3:**
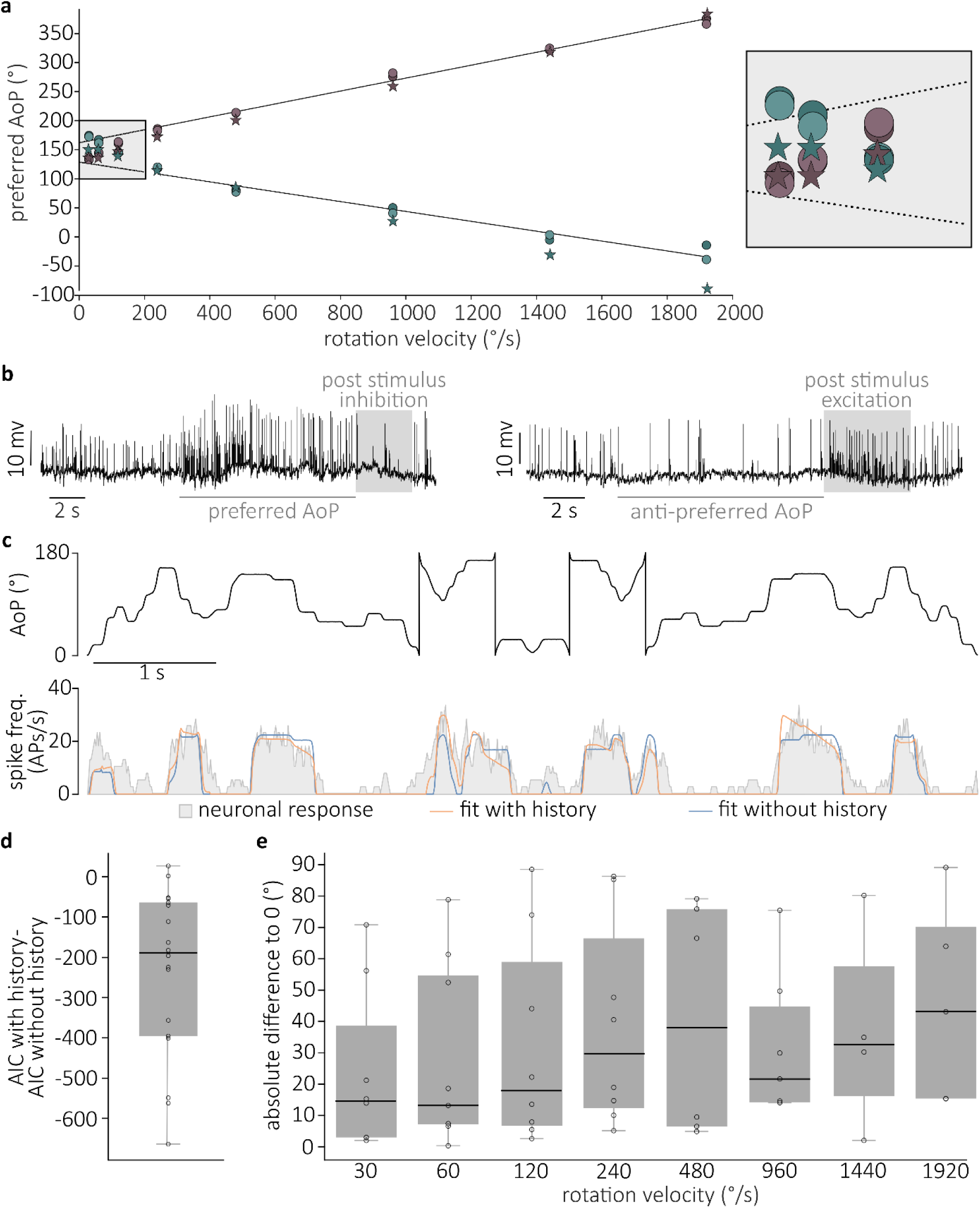
Measured and modeled TL3 neuron responses to continuously rotating and naturalistic stimuli. A) Preferred angle of polarization (AoP) for every velocity and direction (cw: lilac, ccw: turquoise) plotted against rotation velocity. The individual 12 s rotations were divided into two segments of 6 s (first segment, light circles; second segment, dark circles) to illustrate reproducibility of the responses. Circular linear regression analysis (Kempter et al., 2012) was carried out on the five highest rotation velocities. The absolute value of the slope of the regression indicates the delay of the neuron. Dashed line indicates extrapolation of regression towards slow rotation velocities. The stars symbolize the preferred AoP values obtained from the response of the mathematical model using the same stimuli and model parameters as obtained from fit to naturalistic stimulation of the same neuron. The close-up of the low-velocity range on the right shows deviation of the preferred AoPs from the extrapolated regression line (dashed line). Note that ccw values (lilac) in this velocity range are smaller than cw values (turquois), whereas they are bigger at higher velocities. B) Bumblebee CX-neurons show spiking history dependent responses like post-excitatory inhibition (left panel, after presenting preferred AoP for 10 s) and post-inhibitory rebound excitation (right panel, after presenting anti-preferred AoP for 10 s). Recorded data were digitally highpass-filtered using a 0.1 Hz first order Bessel filter. C) Upper panel: naturalistic stimulus sequence, lower panel: recorded neuronal responses (grey, peristimulus time histogram of all 8 stimulus repetitions) and fitted model with spiking history (orange) and without spiking history (blue). D) The difference of Akaike’s information criterion (AIC) for model with history and model without history calculated for each of the recordings (n=18). A value less than 0 indicates that the model with history fits better. E) Comparison of preferred AoP values that were recorded (dots in A), with those calculated from the model parameters that were fitted to the naturalistic stimulus response (stars in A) of the same neurons (n=5, N=10). The absolute difference between these values was calculated and plotted. Boxplots show median, interquartile range (IQR), whiskers with 1.5 x IQR and outliers greater than 1.5 x IQR.

A possible mechanism that may explain the phase-advanced responses at low rotation velocities are spiking-history-effects such as post-excitatory inhibition and post-inhibitory excitation (Figure 3b). In 5 recordings we were able to directly test if polarization-sensitive CX-neurons exhibit such spiking-history-effects. To this end, we calculated the preferred AoP from a cw and a ccw rotation of the polarizer at 60°/s during the recording. Then we presented either the preferred or the anti-preferred AoP for 10 s. Upon termination of the static stimulation with the preferred AoP we observed post-stimulus inhibition whereas the anti-preferred AoP was followed by post-stimulus excitation (Figure 3b).

To test, if phase-advanced firing during continuous slow rotation of the polarizer and the asymmetric responses to the naturalistic-stimulus could indeed be facilitated by spiking-history-effects, we designed two models that simulate spiking activity of polarization-sensitive neurons based on specific stimuli. The first model predicted the spike rate based solely on the AoP of the stimulus and a time delay, while the second model additionally implemented spiking-history-effects (for details see Material and Method and supplementary Figure S1).

To assess which of the models represents the spiking activity of CX neurons more faithfully, we fitted each of them to the measured firing rates in response to the naturalistic stimulus. We used the preferred AoP, the time delay, and the response amplitude, as free parameters in both fits. In the second model, we additionally used the duration of the spiking history that affects the current response, as well as the amplitude of the spiking-history-effects as free parameters (Figure 3c, supplementary Figure S2).

To check which of the two models fitted our data better, we calculated the difference of the Akaike Information Criterion (AIC) values for the model that included spiking history and the one that did not (Figure 3d). Any value of this difference smaller than 0 indicated that the model that included spiking history was favorable. This was the case in 16 out of 18 recordings (88.8%) suggesting that the influence of spiking history can explain the effects we observed in the recordings.

In 5 out of 37 recordings we were able to present both the naturalistic stimulus and the constantly rotating stimuli. This allowed us to test if the same parameter values can be used in the model to describe both the neuronal responses to the naturalistic and to the constantly rotating stimuli. To this end, we first fitted the model to the data obtained during naturalistic stimulation to obtain the specific model parameters for each individual neuron. Using these parameters, we then calculated the response of the model to constantly rotating stimuli with the same angular velocity as the ones tested in the recordings. We then compared the preferred AoPs that resulted from this simulation to the values that were measured during the recordings. An example of this can be seen in Figure 3a, where the circles indicated measured values and the star symbols indicated model data based on the parameters obtained from fitting the model to the naturalistic stimulus. To quantify the agreement between the measured data and the model, we calculated the absolute difference between the preferred AoP (Figure 3e) for each neuron. At lower and medium high rotation velocities (30°/s-240°/s) the median deviation between modelled and measured preferred AoP was between 10.93° and 16.93°, with the smallest value (10.93°) occurring at a rotation velocity of 120°/s. At rotation velocities between 480°/s 1920°/s, the median deviation was between 37.8° and 59.47°.

So far, our results show that spiking activity in CX-neurons in response to polarized-light stimuli depended both on the current stimulus and on spiking history. This means that the spike rate of an individual neuron in response to a specific AoP would be different, depending on its spiking history. Consequently, it would not be possible to infer the AoP of a stimulus from the spike rate of an individual neuron.

To understand how the CX, nevertheless, extracts directional information from the spike rate of polarization-sensitive neurons, we expanded our single-neuron model to a population of neurons. To create a neuronal population with representative model parameters, we averaged the fitted parameters from all recordings of TL3 neurons (n=8; delay, 0.17 s; t_his_, 0.88 s; A, 43.46; A_his_, 0.5; see Methods for model details). TL3 neurons were chosen, because they represented by far the largest group of recordings in our data set. We created a population of 180 neurons with identical parameters, except for the preferred AoP, which was varied between 0° and 179° in 1° increments. We created a 70 s naturalistic stimulus by concatenating the head-orientation data from all tracked bumblebee flights that were published by Boeddeker et al. (2015). The population model was then run both with (Figure 4a) and without the effect of spiking history (Figure 4a’). For better visibility, Figure 4a/a’ show only 20 s out of 70 s (for the full trace see supplementary Figure 3). When including the spiking history, we noticed that the average activity of the population was very variable and depended on the stimulus dynamics. Less dynamic stretches of the stimulus, corresponding to a straighter flight usually led to a decrease in the average spiking activity of the population (Figure 4b and supplementary Figure 4d). This was not the case for the population model without the spiking history. Here, the average population response was always the same, irrespective of the stimulus (Figure 4b’). For the chosen stimulus, the average population activity was slightly smaller for the model which included spiking history (Figure 4b; 6.4 APs/s) than for the model without the spiking history (Figure 4b’; 6.9 APs/s). It should be noted that the flight data from Boeddeker et al. (2015), were obtained around the nest entrance either during turn-back and look behavior or after returning from a foraging trip before entering the nest. It can therefore be expected that during a vector-based outbound or return flight, which is less tortuous, the overall spiking activity of the population is also lower, because of the effects of spiking history on the current spike rate.

**Figure 4:**
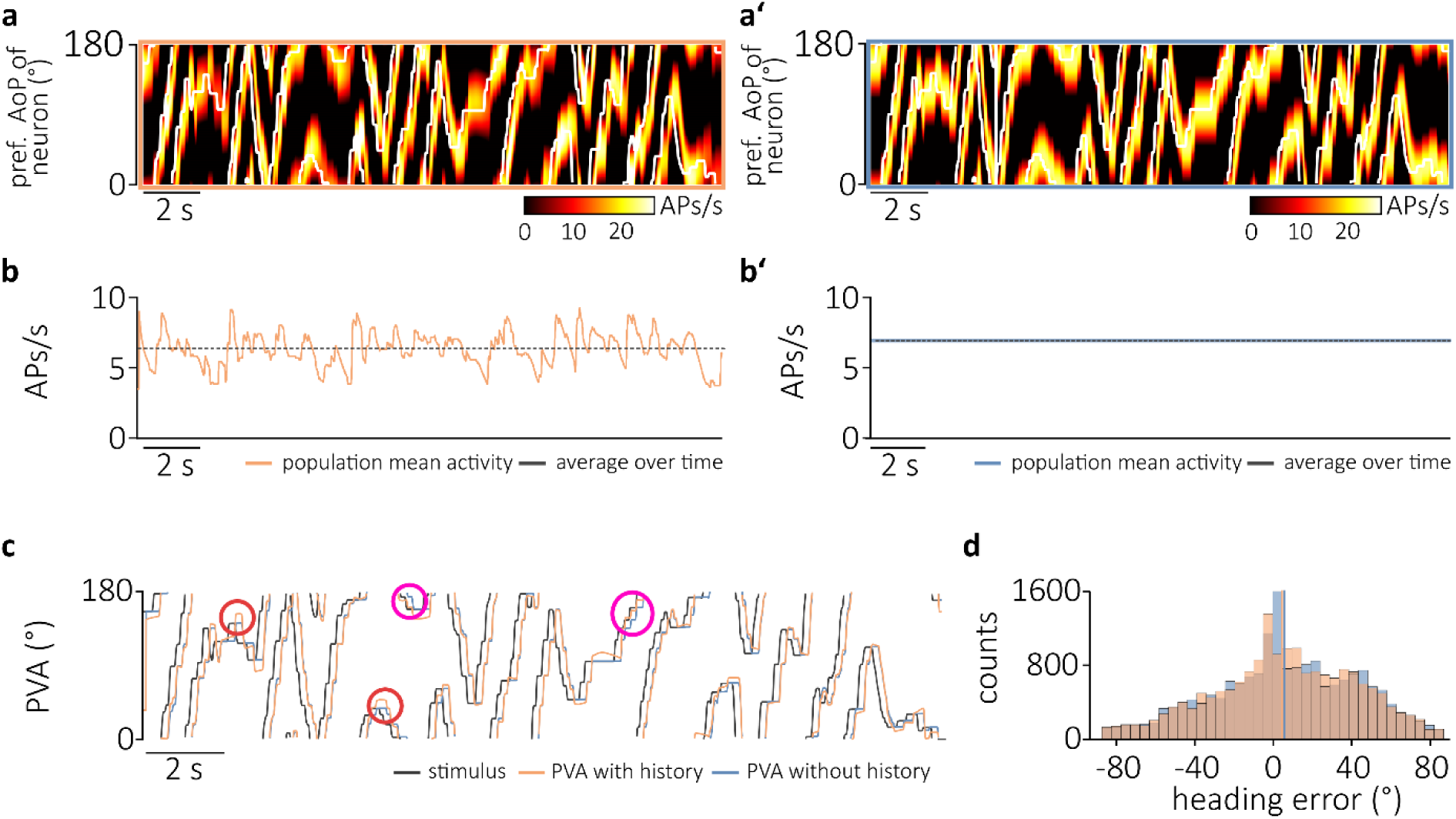
Model of neuronal population responses. The simulation was run using the averaged model parameters from all fitted TL3 neurons n=8 modelling a population of 180 neurons with preferred AoPs between 0° and 179°. A) and A’) Heat maps represent model responses including the spiking history (A) or without spiking history (A’) of the population model to the first 20 s of the stimulus (white) from Boeddeker et al. (2015). The response of each individual neuron in the population corresponds to the response profile along a horizontal line at each preferred AoP. B) and B’) Mean spiking activity of the model population including the spiking history (orange; B) excluding spiking history (blue; B’). Grey line shows average over time. C) Population Vector Average (PVA) with spiking history (orange) or without spiking history (blue). Grey shows the stimulus. Red circles: PVA overshoots the stimulus. Pink circles: PVA from model with history has decreased delay. D) Histogram of heading errors calculated as: heading error = AoP_stimulus_ – PVA. Orange: with spiking history, blue: without spiking history.

To assess, how well the AoP of the stimulus can be extracted from the activity of the population model, we calculated the population vector average (PVA), i.e. the average vector direction that is collectively signaled by the neuronal population. For the model without spiking history, the PVA (Figure 4c; blue line) followed the stimulus (Figure 4c; grey line) perfectly, it was just shifted in time because of the neuronal delays. When spiking history was included in the model (Figure 4c; orange line), the PVA sometimes showed shorter delays (Figure 4c; pink circles) and sometimes overshot the stimulus (Figure 4c; red circles). Nevertheless, when comparing the heading errors, i.e. the difference between the stimulus angle and the PVA, between the two models, the distributions were almost identical (Figure 4d).

To further analyze the population’s response properties, we compared the dynamics of the stimulus to the dynamics of the PVA of both models. Figure 5a shows a part of the stimulus (grey) and the PVAs of the two models (with history: orange; without history: blue), Figure 5b the first derivative of the curves in Figure 5a, i.e. the angular velocities of the stimulus and the PVAs. For easier comparison between the curves, the PVA values (and their derivatives) here were shifted to the left by the delay of the system. For the model without history, the angular velocity profiles of saccadic turns (Figure 5b) were mostly identical to those of the stimulus, whereas this was not the case for the model that included history. For the latter, during many of the saccades, the rate at which the PVA changed (i.e. the PVA velocity) was often larger than the rate at which the stimulus orientation changed (i.e. stimulus velocity). Transferred to a behavioral situation this means, that the rotation velocity that would be signaled by the CX would be larger than the actual rotation velocity of the animal. This seemed to occur particularly often when a cw saccade was followed by a ccw saccade and vice versa, and after longer periods of translational movement. To numerically evaluate this effect, we calculated the ratio between the PVA velocity and the stimulus velocity for each saccade, i.e. values > 1 mean that the PVA changed faster than the stimulus angle during the corresponding saccade. The data were then separated into two groups. Group one included saccades that were preceded by a saccade in the opposite direction, group 2 included saccades that were preceded by a saccade in the same direction. We found that for saccades that were preceded by a saccade in the opposite direction the median ratio between the PVA velocity and the AoP velocity was 1.14 (n = 138), while the median for the other saccades was 1.01 (n = 315). This difference was significant (Mann-Whitney U test, p = 1.3*10^-11^, Figure 5c left). Similarly, during saccades that occurred after at least 200 ms of stationary, i.e. non-rotating AoP, the median ratio between PVA velocity and AoP velocity was significantly elevated to 1.17 (n = 99) compared to 1.02 (n = 354) for the other saccades (Mann-Whitney U test, p = 2.3*10^-11^, Figure 5c, right). Neither of these effects was observed for the model without history, where PVA velocity was always identical to AoP velocity, yielding always a ratio of 1.00 (Figure 5c’). We further evaluated the effect of the duration of stationary AoP before a saccade on the PVA change rate. We found that, for the model with spiking history, the longer the stationary period between two saccades lasted, the bigger the average ratio between PVA velocity and stimulus velocity whereas this had no effect in the model without spiking history (Figure 5d and 5d’). In other words, the neuronal population responded especially fast to angular changes after longer stretches of stationary stimulation. In a behavioral situation this would mean, that the neuronal population becomes more sensitive to angular deviations during straight flight.

**Figure 5:**
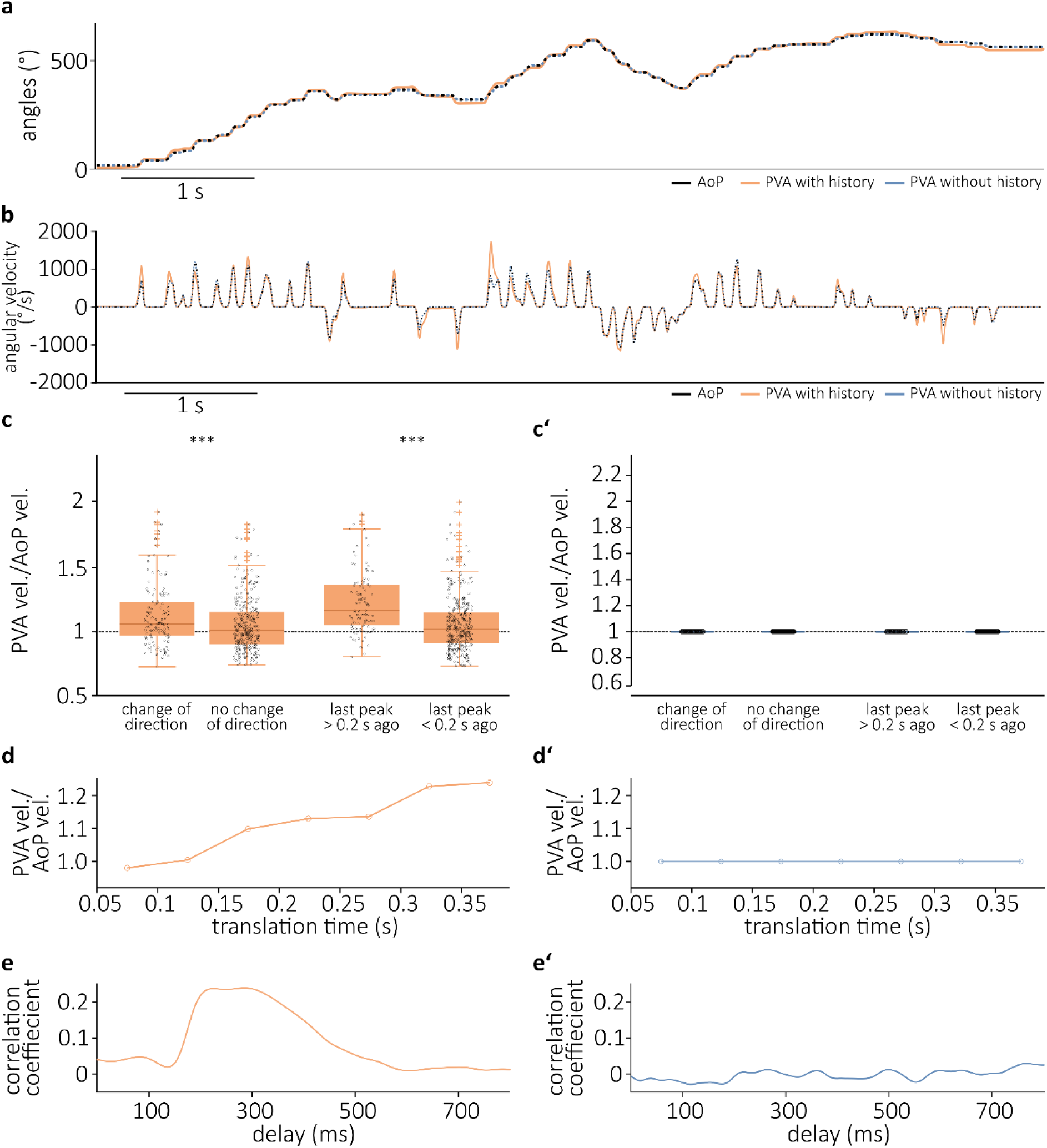
Dynamic properties of neuronal population model. A) Angle of Polarization (AoP; black line), population vector average with history (orange) and without history (blue) for the first 20 s of the naturalistic stimulus (black) created from all flights from Boeddeker et al. (2015). B) First derivative of A), i.e. angular velocity. C) and C’) First and second boxplot: PVA change rate/AoP change rate either after a change of rotation direction (n = 138) or after no change of rotation direction (n = 315). Medians in (C) are significantly different (Mann-Whitney U test, p = 1.3*10^-11^). Third and fourth boxplot: PVA change rate/AoP change rate either after at least 200 ms of translation (n = 99) or after less than 200 ms of translation (n = 354). Medians in (C) are significantly different (Mann-Whitney U test, p = 2.3*10^-11^). Ratio in (C’) is 1 for all data points. Boxplots show median, interquartile range (IQR), whiskers with 1.5 x IQR and outliers greater than 1.5 x IQR. D) and D’) Influence of the duration of translational flight before a saccade on the PVA change rate. This is expressed as the ratio between the peak values of the PVA change rate and the AoP velocity. E) and E’) cross-correlation between mean activity of the population and absolute angular velocity of the AoP (C: with spiking history, C’: without spiking history).

For a flying animal it might be especially important to have a precise knowledge about its heading direction during course corrections, i.e. saccades. Therefore, we wondered if the average spiking rate of the population is related to the dynamics of the stimulus. To test this, we calculated the cross-correlation between the mean activity of the population and the angular velocity of the AoP. The cross-correlation of the population with spiking history (Figure 5e; orange) increased after a delay of roughly 170 ms (the delay implemented in the model), had a plateau between 200 ms and 300 ms and then slowly declined over the next 200 ms. This indicated, that on average saccades were followed by an increase in the population’s activity in the model that included spiking history. No correlation was found for the model without spiking history (Figure 5e’; blue). Since the lowest activity in the population is always 0 spikes/s and the width of the activity “bump” is always the same, a higher mean of the population activity is consistent with a higher signaling contrast.

Taken together our model suggests that polarization-sensitive CX-neurons dynamically adjust their spiking according to spiking history, such that the overall activity of the population decreases during translational flight. Furthermore, spiking-history-effects enable faster signaling of a new heading orientation.

## Discussion

Our new stimulus method allowed us to use naturalistic stimuli to investigate the response dynamics of central complex (CX) neurons during stimulation with polarized light. We show that the responses of these neurons are strongly influenced by the spiking history. As a consequence, the spike rate of an individual neuron is not sufficient to infer the current stimulus. However, using a modelling approach, we show that a population of neurons can reliably encode the angle of polarization (AoP). Furthermore, our model shows that inclusion of spiking history has two beneficial effects on stimulus encoding: 1. Under specific circumstances it allows for faster responses to stimulus changes and 2. it leads to a lower total activity of the population and therefore might enable the system to save energy, especially during straight flight.

The selection of sensory stimuli is crucially important for both physiological and behavioral experiments. Simple, well-defined stimuli are often favored, because they allow to focus on specific aspects of sensory processing by systematically altering just these aspects. Such stimuli can, for example, have sinusoidal characteristics like luminance gratings in visual research. In a more complex form, Gaussian white noise stimuli, i.e. stimuli that contain the same power at all frequencies within a certain frequency band, have been widely used. However, natural visual stimuli are rarely sinusoidal or of Gaussian noise nature, but have specific, non-random statistics, that depend on the habitat and the behavior of the animal (Tolhurst et al., 1992; van Hateren and van der Schaaf, 1996; Dyakova et al., 2019; Nilsson and Smolka, 2021). Studies that used naturalistic stimuli often revealed phenomena that could not be characterized or functionally interpreted with artificial stimuli (Rieke et al., 1995; Felsen and Dan, 2005; Pfeiffer and French, 2015; Dyakova and Nordström, 2017). Likewise, naturalistic stimulation in this study allowed us to quantify and functionally interpret the spiking history dependence of CX-neurons, and to estimate the population responses to behaviorally relevant stimuli.

A particularly interesting aspect of our data were the systematic shifts of their tuning to the AoP depending on rotation velocity and direction. While the shift at high velocities was expected and is a consequence of delays in the system, the preferred AoP at low rotation velocities was shifted in the opposite direction i.e. against the rotation direction. Although this phenomenon has been briefly mentioned in the literature before (Sakura et al., 2008), it has not been systematically investigated. Interestingly a similar phenomenon has been described in the primary visual cortex of vertebrates. Orientation selective neurons in V1 change their preferred orientation tuning towards the orientation of an adapting stimulus if it is presented for a longer duration, while they change it away from it during shorter presentation (Ghisovan et al., 2009). It is assumed, that this so called “attractive shift” facilitates orientation discrimination around the area of the adapting stimulus. This interpretation is consistent with our data. Longer periods of stationary polarized light stimuli lead to both stronger and faster responses when the stimulus changed.

What are the possible mechanisms underlying the spiking-history-effects we observed? There are at least three mechanisms that might explain our observations: 1. Adaptation at the level of the polarization-sensitive dorsal rim area (DRA) of the compound eye, 2. Neuronal adaption within the polarization-vision pathway, and 3. Effects of neuronal plasticity within the network. These possibilities are not mutually exclusive, and it is possible that all three contribute to the observed effects. The DRA of insects is organized such that individual ommatidia are specialized to sense two orthogonal AoP. Across the entire DRA, the preferred AoP systematically shifts, such that all possible orientations are covered. Therefore, during slow rotation or stationary presentation of the polarized-light stimulus, the photoreceptors that align with the current AoP will inevitably adapt while those that are oriented perpendicularly will recover from adaptation, thus creating an imbalance in the inputs to the polarization pathway. This likely shifts the percept of a polarized-light stimulus. While slowly rotating through the anti-preferred AoP of the neuron, photoreceptors that are tuned to this angle will become more adapted, while those that are tuned to the perpendicular angle will become less adapted. As a result, when rotating towards the preferred AoP of a neuron, its peak activity will be reached faster. The second possibility, adaptation at the neuronal level, could happen, because TL neurons in insects exist as a set of cells that are each tuned to different angles of polarization. Similar to the receptor layer, a stationary stimulus could, depending on its AoP, adapt individual neurons of the population of TL neurons, or neurons along the polarization pathway (Pfeiffer and Kinoshita, 2012), while others recover from previous adaptation. TL neurons in many insect species respond both to polarized light and a simulated sun stimulus (monarch butterfly: Heinze and Reppert, 2011, dung beetle: el Jundi et al., 2015, locust: Pegel et al., 2018). Since these two stimuli are perceived by different parts of the compound eye, future experiments could use a simulated sun stimulus to disentangle possibility 1 and 2. If adaptation happens mainly in the neurons, it should be strongly dependent on the dynamics of the stimulus and less dependent on the nature of the stimulus (unpolarized or polarized light). If adaptation in the receptors (or their synapses) is more relevant, bigger differences should be seen between unpolarized and polarized-light stimulation. The third possibility, i.e. changes within the compass network, could happen at any stage of the polarization vision pathway. However, studies in *Drosophila* show, that all R-neurons (the *Drosophila* homologues of TL neurons) of a specific type are connected to all other individual neurons of the same type, as well as to all EPG neurons, which are regarded as head-direction neurons (Seelig and Jayaraman, 2015; Hulse et al., 2021). It has further been demonstrated that this connection is highly plastic and can establish a heading representation that may be anchored to different visual environments within minutes (Fisher et al., 2019; Kim et al., 2019). It is conceivable, that this network is also capable of short-term plastic changes that contribute to the observed effects.

The ability of the central complex neurons to dynamically adjust their properties according to the spiking history allows for a more flexible coding. Because phototransduction and neuronal transmission require time, compass neurons can never signal the current heading direction instantaneously. The overall error of the heading signal, which arises from this delay does not differ between the model that implements spiking history and the one that does not, at least for the stimulus sequence that was used in this study. However, under specific circumstances, such as longer durations of stationary stimulation, or switches between clockwise and counter-clockwise rotations, the angular heading signal of the population model changed faster than the stimulus angle. In a behavioral context, this could be used to anticipate future head directions as has been suggested for some head direction cells in rats (Blair and Sharp, 1995). Such an anticipatory signaling could be used to allow the bees to induce the end of a saccadic sequence before a desired heading is reached to compensate for their moment of inertia. The mechanism could also be an important factor in shaping the capabilities of bumblebees to maintain a straight course: During translational flight the system would become specifically responsive to deviations from the current heading direction and therefore allow for quicker course corrections.

Another advantage of the spiking history dependent properties of central complex neurons is that during straight flight, the overall spiking activity of the neuronal population is strongly decreased, therefore reducing the energy consumption of the system. It is well documented, that the nervous system consumes considerable amounts of energy and that energy consumption is a driver in the evolution of sensory systems (Niven and Laughlin, 2008). The naturalistic stimuli we used to probe the CX-neurons during the recordings were derived from bumblebees that were either leaving the hive or returning to it (Boeddeker et al., 2015). In these situations, the animals perform stereotypical behaviors to either memorize or find the entrance of the hive. These flight patterns are usually rather tortuous containing many saccades, which made them particularly useful as a stimulus to probe the dynamic properties of the neurons. However, a considerable fraction of foraging, consists of shuttling between the hive and a food patch - a part of the flight which is less tortuous. In our model, stationary stimuli, reduced the average spiking activity of the neuronal population almost by a factor of 3 (Supplemental Figure 4), suggesting that during straight flight, the entire neuronal population reduces its spiking activity and thus conserves energy.

Taken together, our study shows that populations of polarization-sensitive CX-neurons in the brain of the bumblebee are able to signal heading direction based on naturalistic inputs, with responses that strongly depend on spiking history. Moreover, the dynamic properties of the neurons lead to a decrease in overall activity during periods of straight flight and allow for a faster adjustment of directional coding.

## Acknowledgments

We are grateful to Dr. Barbara Webb for helpful discussion about the model and for pointing us to the Akaike Information Criterion, and to Dr. James J Foster for help implementing it and advice on fitting, to Drs. Christine el Jundi and Frederik Zittrell for providing code for the circular-linear correlation analysis, and to Katja Tschirner and Evelyn Beetz for animal care. The study was supported by the German Research Foundation (Deutsche Forschungsgemeinschaft, DFG) equipment grant INST93/829-1, and research grant PF714/5-1.

## Supplementary information

**Supplemental Figure 1:**
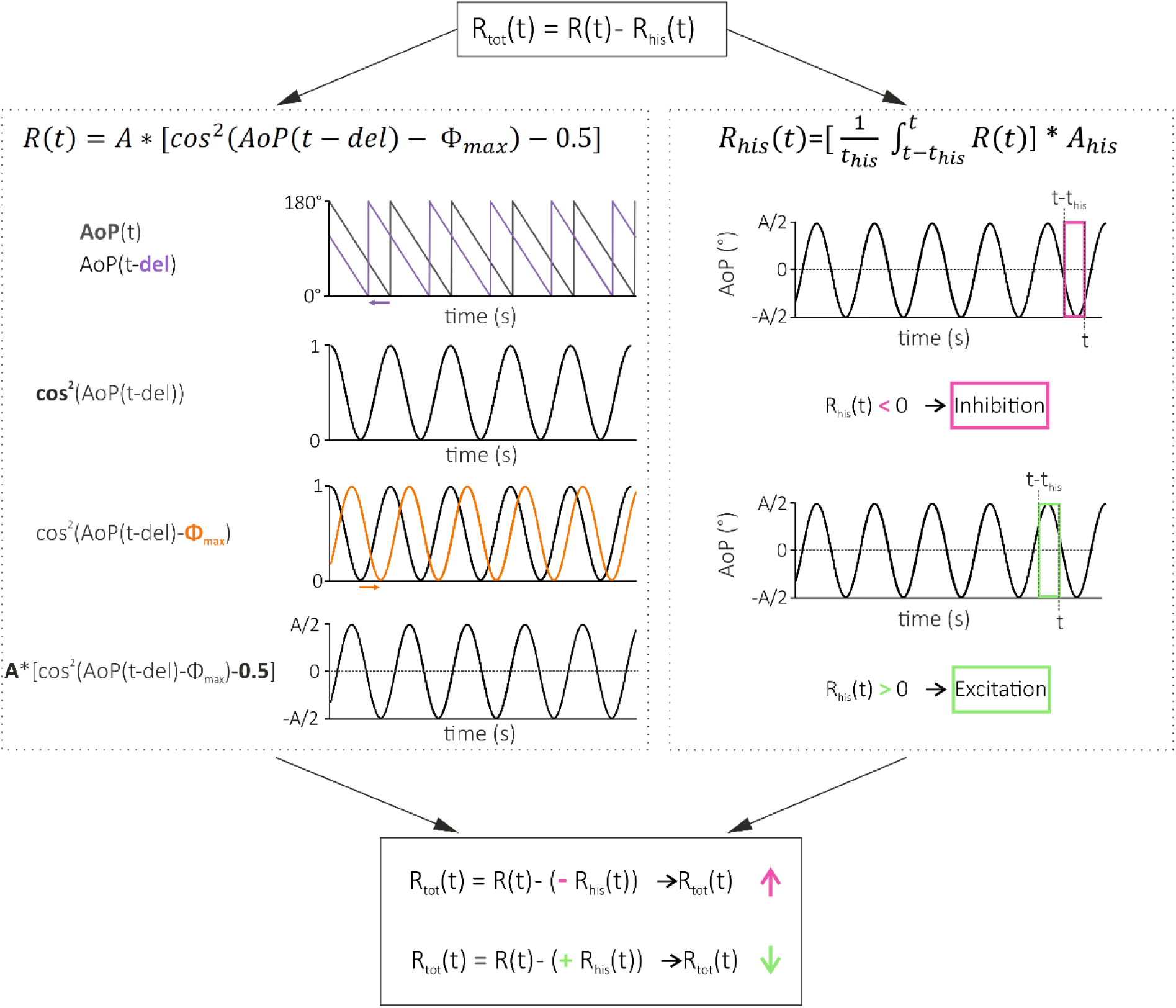
Graphical illustration of model calculations. Neuronal firing rate *(R)* over time *(t)*, elicited by the stimulus was modeled as a cos^2^ function of the angle of polarization (AoP). Where *A* is the response amplitude, *del* is the system’s delay and *Φ_max_* is the preferred angle of polarization of the neuron. To accommodate for spiking-history-effects, a spiking history *(R_his_)* was implemented into the model. Spiking history was defined by two parameters: the length of a rectangular window, over which past responses are averaged and *A_his_,* a scaling factor for the spiking-history-effect. *R_his_* was then subtracted from the response to the current stimulus *(R)*, to give the total response *R_tot_*. Since the model calculates spike rate, negative values were replaced by zeros.

**Supplemental Figure 2:**
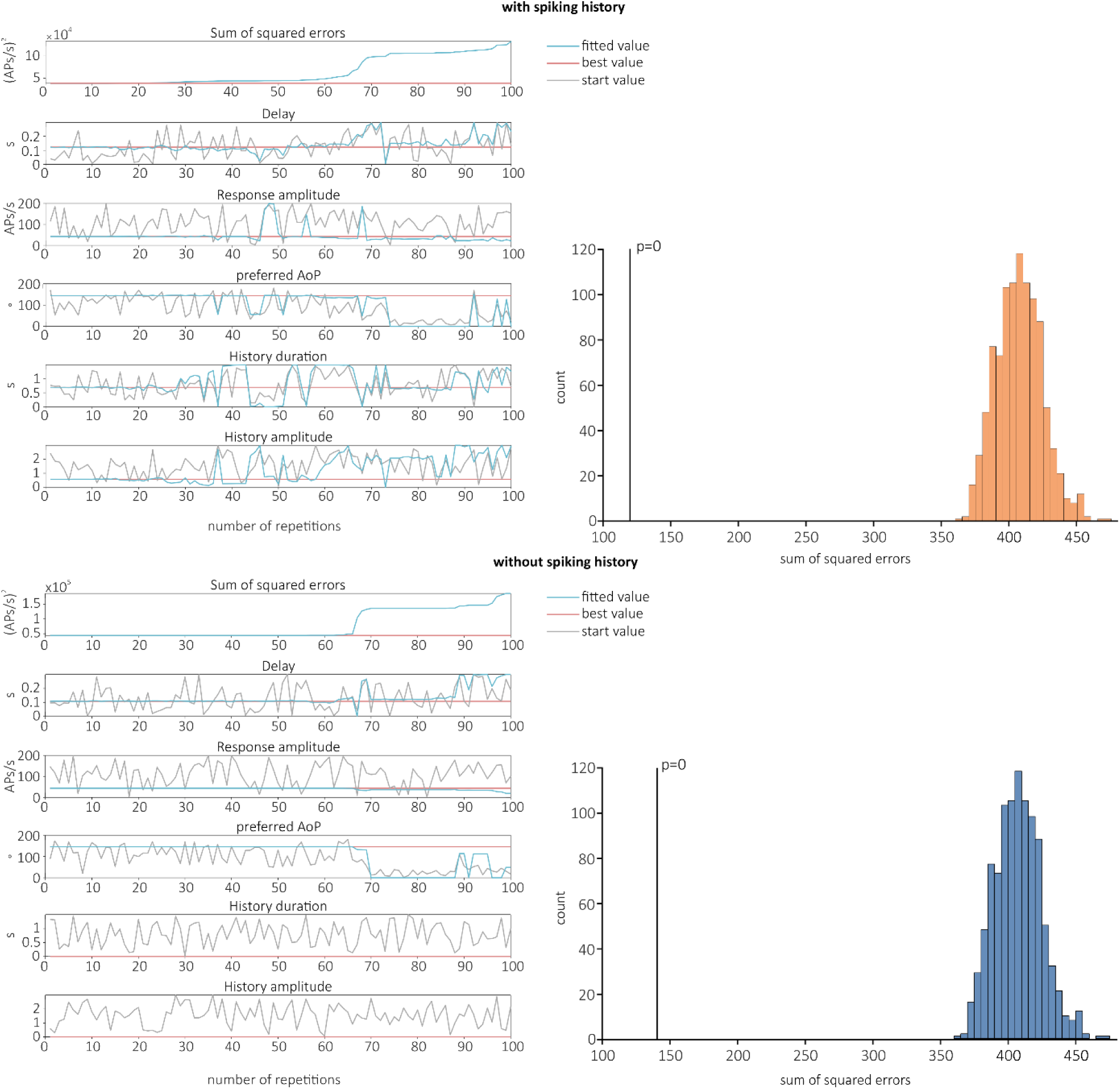
Starting parameter and Bootstrapping for model fit. The parameters preferred AoP, delay, response amplitude, history duration (length of memory window), and amplitude of spiking history were used as free parameters. To avoid the detection of local minima in the fitting procedure, we repeated the fitting procedure 50 times for each recording and initialized the fitting parameters each time with random start values. The fitted parameter values were then obtained from the best of the 50 fits. To assess the quality of the fit, we used a bootstrapping method. We repeated the fitting procedure 1000 times, but instead of fitting the model to the original PSTHs, we randomly permuted the bins of the PSTHs before the fit. We then compared the sum of squared errors of the fit to the actual data to the sum of squared errors of the permuted data.

**Supplemental Figure 3:**
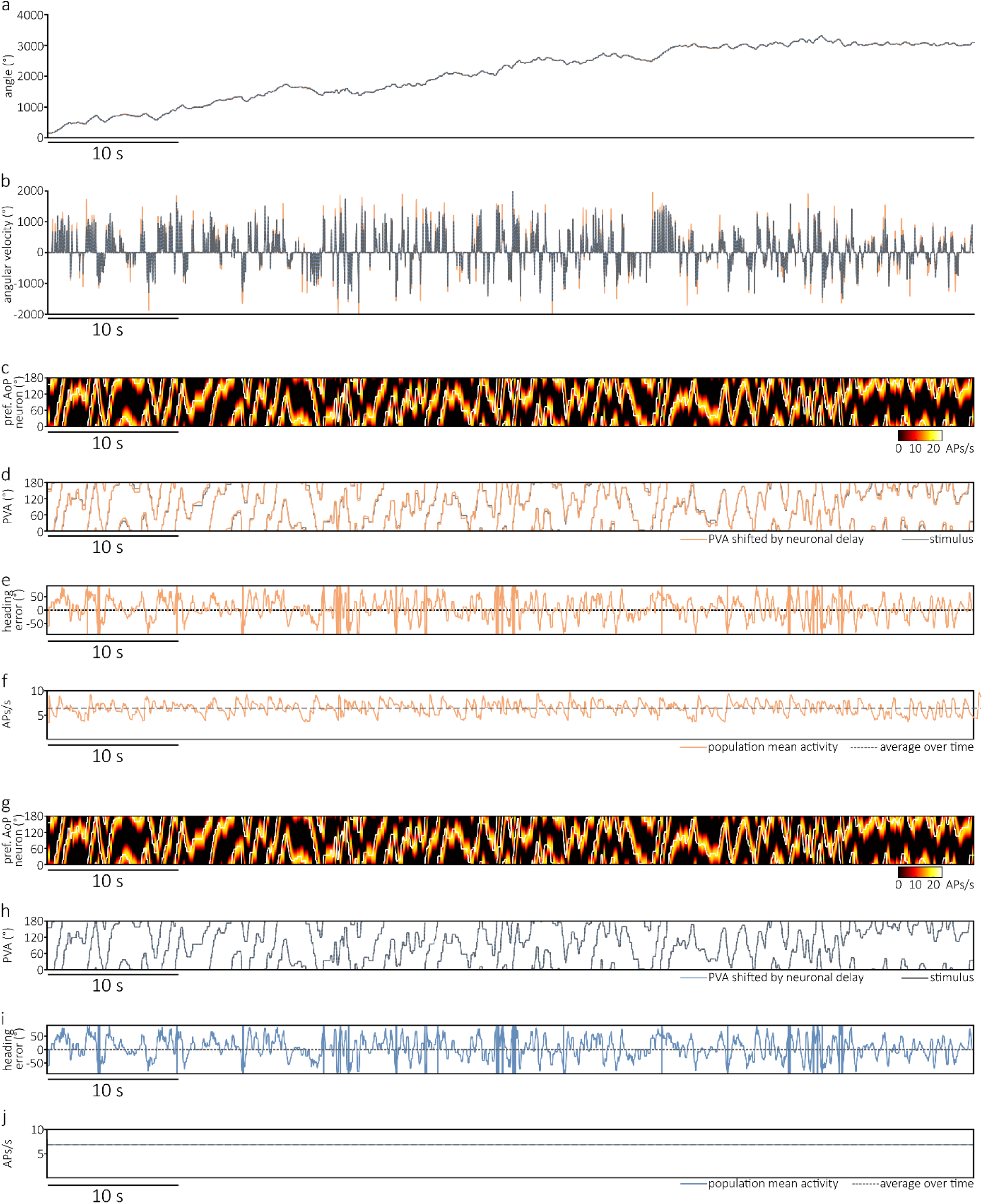
Results for concatenated flight tracks of all recorded bumblebee flights from Boeddeker et al. 2015. A) grey: Angle of Polarization (AoP), orange: Population vector average (PVA) for model with spiking history, blue: Population vector average (PVA) for model without spiking history. B) First derivative from the track from A showing angular velocities. The PVA values and their derivatives are shifted by the delay of the system for better comparison to the stimulus. C) The heatmap represents the response of the modelled population of neurons (with spike history) to the stimulus (white line). D) Grey shows the stimulus, orange the population vector average (PVA) shifted to the left by the neuronal delay and calculated from the responses in C. E) Heading Error calculated as AoP – PVA. F) Orange: Population mean activity from the responses including models spiking history, grey: averaged activity over time. G) The heatmap represents the responses for population of neurons to the stimulus (white line) without the models spiking history. H) Grey shows the stimulus, blue the population vector average (PVA) shifted to the left by the neuronal delay and calculated from the responses in G. I) Heading Error calculated as AoP – PVA. J) Blue: Population mean activity from the responses including models spiking history, grey: averaged activity over time.

**Supplemental Figure 4:**
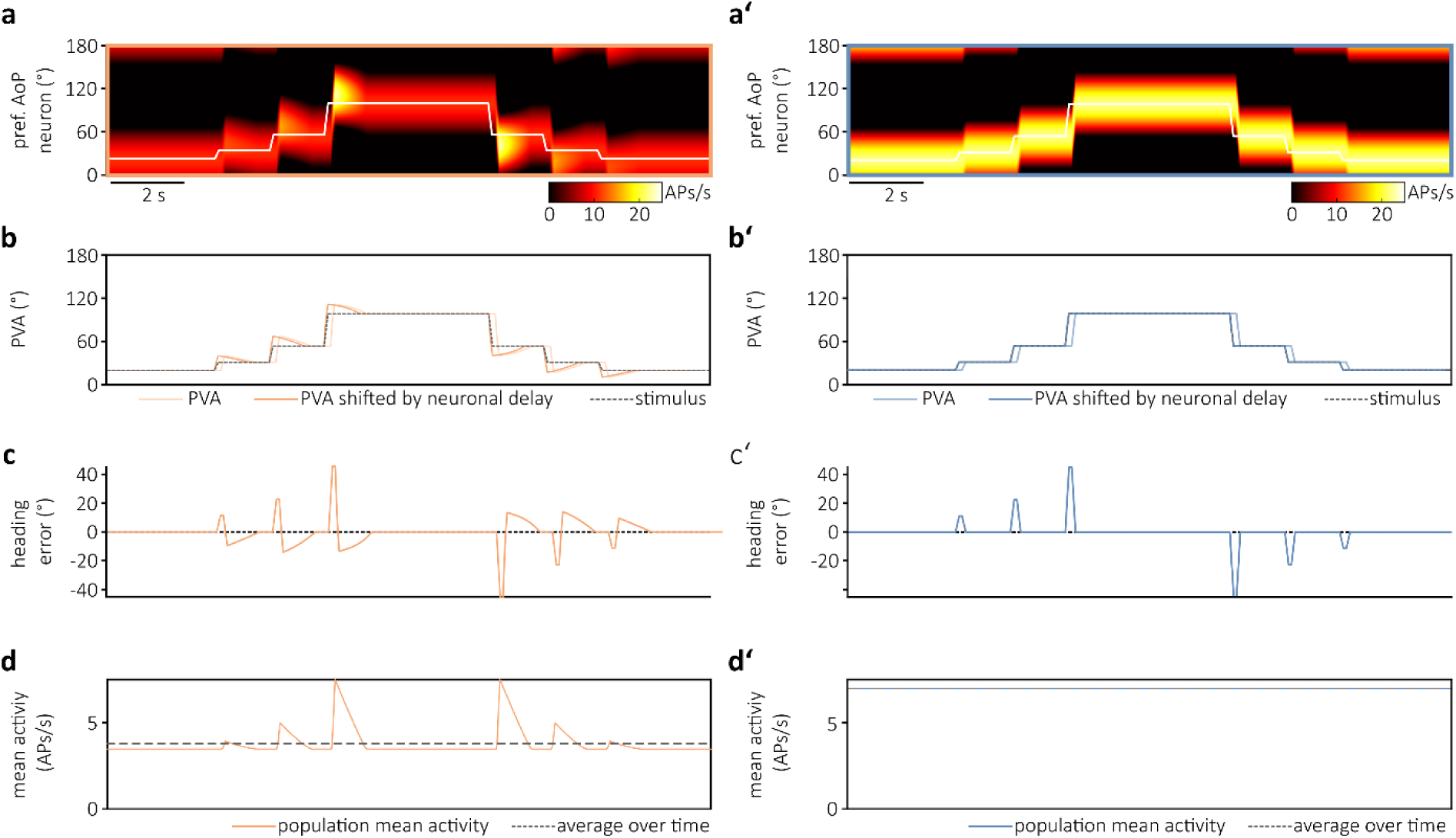
Population model response to stimulus with sparse rotations. A) and A’) The heatmaps represent the responses for a modelled population of neurons to an artificial stimulus with sparse rotations (white line) including the models spiking history (A) or without the spiking history (A’). Note that population activity (with history) declines over time and is higher, the larger the rotation angle. B) Grey shows the stimulus, orange the population vector average (PVA, light orange) shifted to the left by the neuronal delay (dark orange) and calculated from the responses in A. B’) Grey shows the stimulus, blue the population vector average (PVA, light blue) shifted to the left by the neuronal delay (dark blue) and calculated from the responses in A. C) and C’) Heading Error calculated as AoP – PVA, in orange for model with (orange) and without (orange) spiking history. D) and D’) Population mean activity from the responses including models spiking history (orange) or without spiking history (blue), grey: averaged activity over time.

## Notes

### Competing Interest Statement

The authors have declared no competing interest.

